# A Cell Autonomous Free fatty acid receptor 4 - ChemR23 Signaling Cascade Protects Cardiac Myocytes from Ischemic Injury

**DOI:** 10.1101/2024.11.26.625260

**Authors:** Sara J. Puccini, Chastity L. Healy, Brian A. Harsch, Ahmed R. Ahmed, Gregory C. Shearer, Timothy D. O’Connell

## Abstract

Acute myocardial infarction (AMI) causes ischemic damage and cardiac remodeling that ultimately progresses into ischemic cardiomyopathy (ICM). Coronary revascularization reduces morbidity and mortality from an MI, however, reperfusion also induces oxidative stress that drives cardiac myocyte (CM) dysfunction and ICM. Oxidative stress in CMs leads to reactive oxygen species (ROS) production and mitochondrial damage. Free fatty acid receptor 4 (Ffar4) is a GPCR for long chain fatty acids (FA) that is expressed in multiple cell types including CMs. We have recently shown that CM-specific overexpression of Ffar4 protects the heart from systolic dysfunction in the context of ischemic injury. Mechanistically, in CMs, Ffar4 increases the levels of 18-hydroxyeicosapentaenoic acid (18-HEPE), an eicosapentaenoic acid (EPA)-derived, cardioprotective oxylipin (oxidatively modified FA). 18-HEPE is the precursor for resolvin E1 (RvE1), a cardioprotective, specialized pro-resolving mediator (SPM) that activates the GPCR ChemR23. We hypothesize Ffar4 in CMs protects the heart from oxidative stress and ischemic injury through activation of a CM-autonomous, Ffar4-ChemR23 cardioprotective signaling pathway. Here, we developed an *in vitro* hypoxia reoxygenation (H/R) model (3 hours of hypoxia, 17 hours of reoxygenation) in adult CMs as a model for ischemic injury. In adult CMs subjected to H/R, TUG-891, an Ffar4 agonist, attenuated ROS generation and TUG-891, 18-HEPE, and RvE1 protected CMs from H/R-induced cell death. More importantly, we found that the ChemR23 antagonist α-NETA prevented TUG-891 cytoprotection in adult CMs subjected to H/R, demonstrating that ChemR23 is required for Ffar4 cardioprotection. In summary, our data demonstrate co-expression of Ffar4 and ChemR23 in the same CM, that Ffar4, 18-HEPE, and RvE1 attenuate H/R-induced CM death, and that ChemR23 is required for Ffar4 cardioprotection in H/R support a CM-autonomous Ffar4-ChemR23 cardioprotective signaling pathway.

## Introduction

According to the American Heart Association 9.3 million people, 3.2% of the US population, have had a myocardial infarction (MI) (1). While coronary revascularization has decreased mortality following an acute MI, the incidence of ischemic cardiomyopathy (ICM) and heart failure (HF) post-MI has continued to rise (2).

Revascularization post-MI can lead to severe cardiac myocyte (CM) dysfunction largely driven by oxidative stress leading to mitochondrial damage and cell death, provoking a multiphasic inflammatory response to drive ICM (3). Therapeutic management of patients post-MI primarily targets reducing afterload and minimizing the risk of a future MI (4). Currently, there are no FDA-approved therapeutics that specifically target reducing oxidative stress post-MI.

Free fatty acid receptor 4 (Ffar4) is a Gq-coupled G-protein coupled receptor (GPCR) for long chain saturated, monounsaturated, and polyunsaturated fatty acids (SFA, MUFA, and PUFA). Previously, we have demonstrated that loss of Ffar4 worsens cardiac outcomes in multiple experimental models of HF, including pressure overload (5), cardiometabolic disease (6), and ischemia-reperfusion (I/R) injury (7). Specifically, in response to I/R, loss of Ffar4 worsened left ventricular (LV) systolic function, without an effect on infarct size in male and female mice, whereas CM-specific overexpression of Ffar4 attenuated LV systolic dysfunction post-I/R (7). Generally, we have found that the loss of Ffar4 impairs cardiac function without an effect on fibrosis.

In CMs, we have demonstrated that Ffar4 signaling through cytoplasmic phospholipase A2α (cPLA2α) induces the production of the eicosapentaenoic acid (EPA, an ω3-PUFA)-derived oxylipin, 18-hydoxyeicosapentaenoic acid (5). Further, loss of Ffar4 was associated with reductions in 18-HEPE levels in the heart and HDL and correlated with worsened outcomes in pathologic mouse models of HF (5, 6).

Interestingly, a prior study found that 18-HEPE attenuated pathologic remodeling in pressure overload (8). 18-HEPE is the precursor for resolvin E1 (RvE1), a cardioprotective, specialized pro-resolving mediator (SPM) that activates the GPCR ChemR23. Prior studies also indicate that RvE1 attenuates cardiac ischemic injury *ex vivo* (9) and *in vivo* (10, 11). Together, these data suggest that Ffar4-mediated production of EPA-derived oxylipins is cardioprotective.

Here, we propose a CM-autonomous Ffar4-ChemR23 cardioprotective signaling pathway that protects CMs from oxidative stress. Specifically, we hypothesize that Ffar4 induces the production of 18-HEPE, which is converted to RvE1 to activate ChemR23, activation of which protects CMs from oxidative stress as illustrated in **Figure 1**. Briefly, our results indicate that Ffar4 and ChemR23 are co-expressed in the same CM, a prerequisite for our model. Using a model of hypoxia-reoxygenation (H/R) to model I/R *in vitro*, we found that activation of Ffar4 attenuated H/R-induced oxidative stress and CM death. Similar results were observed with 18-HEPE and RvE1. Finally, we found that the ChemR23 antagonist, α-NETA, ablated Ffar4 cardioprotection in H/R, indicating ChemR23 is required for Ffar4 cardioprotection. In summary, we have identified a novel CM-autonomous Ffar4-ChemR23 cardioprotective signaling pathway that attenuates oxidative stress.

**Fig. 1:**
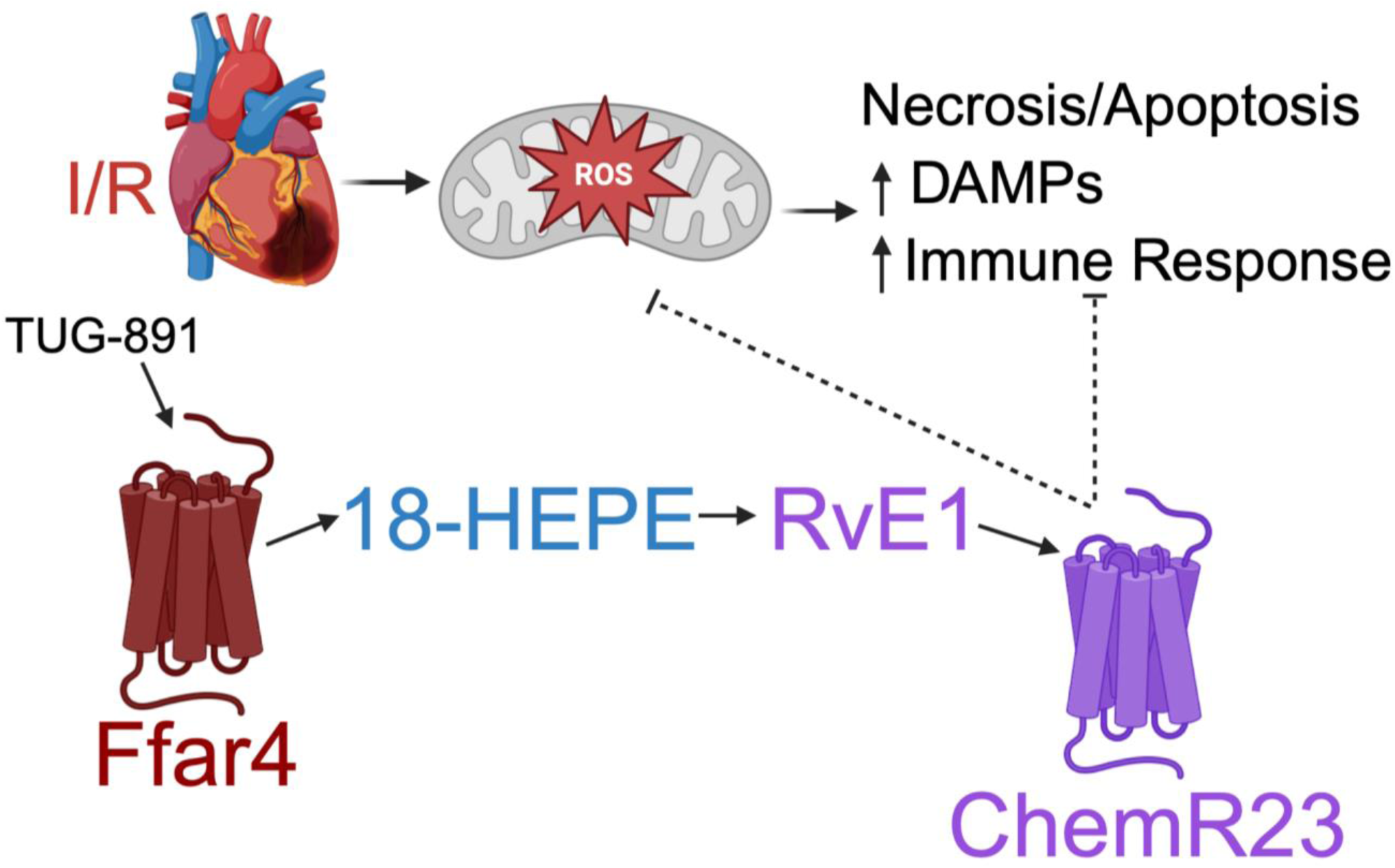
A cardiac myocyte (CM) autonomous Ffar4-18-HEPE-RvE1-ChemR23 signaling pathway attenuates ischemic injury. We propose that in adult CMs, activation of Ffar4 induces production of the eicosapentaenoic acid (EPA)-derived, cardioprotective oxylipin 18-hydroxyeicosapentaenoic acid (18-HEPE). Subsequently, 18-HEPE is further oxidatively modified to produce the specialized pro-resolving mediator (SPM) resolvin E1 (RvE1). RvE1 is an endogenous agonist for ChemR23, and activation of ChemR23 attenuates oxidative stress to prevent ischemic injury in adult CMs.

## Methods

### Isolation and culture of adult cardiac myocytes

We previously published a comprehensive step-by-step procedure for the isolation and culture of adult mouse cardiac myocytes that was used in this study (12).

### Single CM Real Time Quantitative PCR (RT-qPCR)

Adapting a previously described protocol (13), freshly isolated adult CMs in standard perfusion buffer were individually pipetted into an single PCR tube in ∼5 μl volume using a Stripper Pipette (MXL3-STR) equipped with a 100 µm tip (MXL3-100) from Cooper surgical (Brand: ORIGIO) and flash-frozen in liquid nitrogen.

#### Reverse Transcription

cDNA libraries were synthesized from single CMs in a two-step process in a final total volume of 20 μL. Step 1, the following reagents were added to each tube containing 1 cell in ∼5 μL: 2 μL M-MuLV Reverse Transcriptase Reaction Buffer (New England Biolabs (NEB), Ipswich, MA, Cat. #M0253), 5 μM Oligo(dt) Primer (ThermoFisher, Waltham, MA, Cat. #18418012), 60 μM Random Hexamer Primer (NEB, Cat. #S1330S), and Nuclease Free Water up to a final volume of 16 μL. The tubes were heated to 65°C for 10 minutes then placed on ice. Step 2, the following reagents were added to each reaction: 1 mM dNTP (NEB, Cat. #N0447), 20 U RNase Inhibitor (NEB, Cat. #M0353), 35 U Reverse Transcriptase (NEB, Cat. # M0253), and nuclease free water up to final volume of 20 μL. In a thermocycler, the reactions were heated to 25°C for 10 minutes, 55°C for 30 minutes, and then 85°C for 5 minutes, and the reactions were subsequently cooled on ice. cDNA preparations were then purified utilizing a Qiagen QIAquick PCR purification kit (Qiagen, Germantown, MD) with an elution volume of 50 μL.

#### Pre-Amplification

Primers for specific genes of interest were pooled based on annealing temperatures and level of expression for “primer pool” pre-amplifications.

Pool 1: RPL32 and cTNT, Pool 2: Ffar4, and Pool 3: ChemR23. Primer pool mixes were prepared at a concentration of 1 μM. For each pre-amplification reaction, we combined 2.25 μL of the 1 μM primer pool mix, 10 μL 2x iTaq Universal SYBR Green (BioRad, Hercules, CA, Cat. #1725120), 5 μL of cDNA, and 2.75 μL nuclease free water for a final volume of 20 μL. Reactions were amplified using the following cycling parameters: 1X: 95°C for 3 minutes; 19X: 95°C for 20 seconds, annealing temp for specific primers for 4 minutes, 72°C for 20 seconds; and 1X: 72°C for 5 minutes, and following the reaction, amplified cDNA was diluted 1:2 with nuclease free water.

#### RT-qPCR

RT-qPCR was performed using a BioRad CFX96 Real-Time System thermocycler. For each reaction we combined 1.5 μL of cDNA, 100nM primers, 5 μL 2x iTaq Universal SYBR Green (BioRad, Cat. #1725120) and nuclease free water up to 10 μL. Reactions were amplified using the following cycling parameters: 1X: 95°C for 30 seconds and 39X: 95°C for 5 seconds, annealing temp for 30 seconds + data capture. All qPCR results are verified with DNA gel electrophoresis.

**Table 1:**
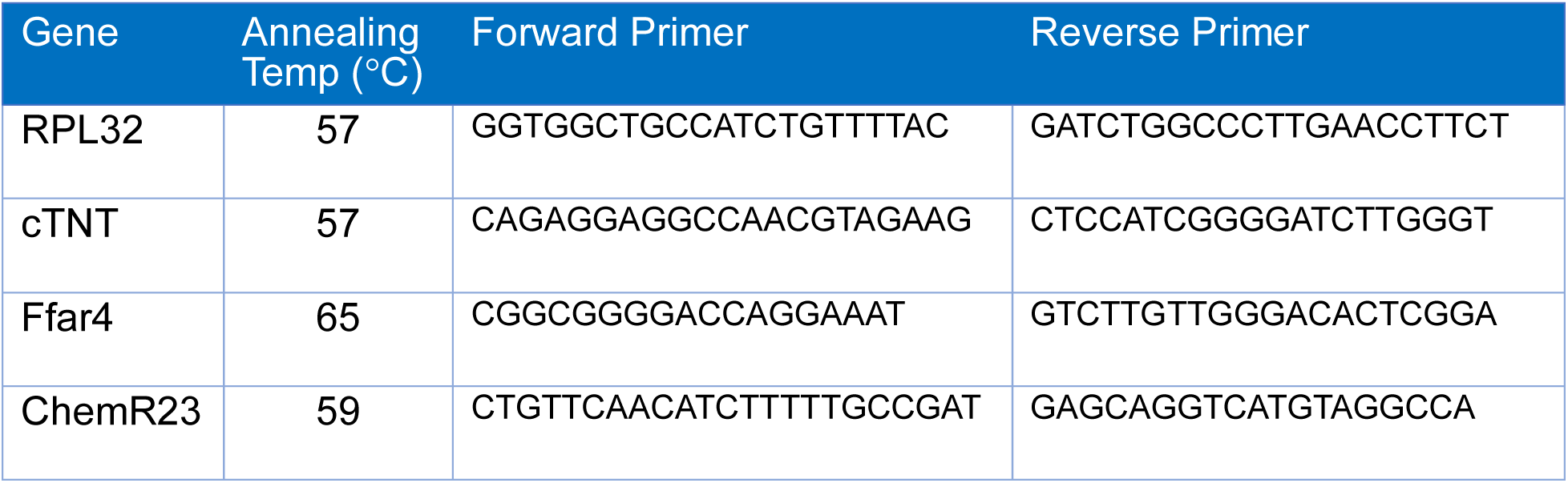
RT-qPCR primer sequences.

### Hypoxia/Reoxygenation (H/R)

Adult CMs were isolated and cultured in 35 mm tissue culture dishes as described above and subjected to H/R using an adaptation of a previously described protocol (14). Briefly, prior to hypoxia, CMs are pretreated with either TUG-891 (10 µM, overnight, Cayman Chemicals, Ann Arbor, MI, Cat. #17035), 18-HEPE (100 nM, 2 hours, Cayman Chemicals, Cat. #32840), resolvin E1 (100 nM, 2 hours, Cayman Chemicals, Cat. #10007848), α-NETA (100 nM, overnight, Cayman Chemicals, Cat. #15125), or vehicle. To prepare hypoxic culture medium prior to adding to CMs, culture medium was bubbled in a 50ml conical for 30 minutes with a nitrogen gas mixture containing 94% N_2_, 5% CO_2_, and 1% oxygen. To induce hypoxia, the cell culture medium was changed to hypoxic medium with either the treatment of interest or vehicle. CMs were then placed inside a hypoxia chamber (STEM CELL Technologies, Vancouver, BC, Canada, Cat. #27310) filled with the nitrogen gas mixture for 3 hours.

After 3 hours, reoxygenation was initiated by removing CMs from the hypoxia chamber and changing the culture medium back to normoxic medium containing the treatment of interest or vehicle for up to 17 hours. CM death was measured at 17 hours, whereas CellROX staining to measure ROS generation was measured after 4 hours of reoxygenation.

### Quantification of CM death induced by H/R

Following 17 hours of reoxygenation, cell death was quantified by assessing cell morphology (rod shape: viable; round: dead) for duplicate culture dishes for each condition, quantifying at least 100 myocytes from 5 randomly selected fields in each culture dish as we have previously described (15–18).

### Quantification of Reactive Oxygen Species (ROS) induced by HR

Following 4 hours of reoxygenation, CM ROS was quantified using CellROX Deep Red (ThermoFisher, Cat. #C10422), a fluorescent dye used to detect ROS generation. Briefly, CellROX (5 μM) was added to the culture medium for 30 minutes at 37°C. After 30 minutes, CMs were fixed in a two-step process, 1) 10 minutes with 2% paraformaldehyde, and 2) 10 minutes with 4% paraformaldehyde. After fixation CMs were stained with DAPI (4′,6-diamidino-2-phenylindole) and mounted on microscope slides. To detect CellROX fluorescence, CMs were excited at 649 nM and images were captured using a 667 nM filter with a 10X objective on a Keyence BZ-X810 Fluorescence Microscope (Keyence, Osaka, Japan). Mean fluorescent intensity (MFI) for individual CMs was measured using ImageJ (19), and background intensity was subtracted.

### Oxylipin Production induced by H/R

#### CM preparation

Following 17 hours of reoxygenation, cell culture medium was removed, CMs were rinsed with warm PBS, and placed in a 1:1 mixture of PBS:Methanol. CM were scraped and collected into the PBS:Methanol solution and flash-frozen in liquid nitrogen.

#### Sample Extraction

Samples were thawed on ice and extracted using a previously well-defined protocol (5). CMs were spiked with BHT/EDTA (0.2 mg/mL), a surrogate containing deuterated oxylipins (20 uL of 1000 nM concentration), and subjected to liquid-liquid extraction to isolate lipid content. Samples were split in two, a non-hydrolyzed portion for free oxylipin measurement, and a hydrolyzed portion for total oxylipin measurement using 0.1 M methanolic sodium hydroxide and further purification by solid phase extraction using a Chromabond HLB sorbent column. Oxylipins were eluted with 0.5 mL of methanol with 0.1% acetic acid and 1 mL of ethyl acetate, dried under nitrogen stream, and reconstituted with 100 nM 1-cyclohexyluriedo-3-dodecanoic acid (CUDA) in methanol acetonitrile (1:1).

#### LCMS Analysis

Samples were analyzed by liquid chromatography using a Waters Acquity UPLC coupled to Waters Xevo triple quadrupole mass spectrometer equipped with electrospray ionization source (Waters, Milford, MA). A volume of 5 μL was injected and separation was performed using a CORTECS UPLC C18 2.1 x 100 mm with 1.6 μM particle size column for oxylipin analysis. Flow rate was set at 0.5 mL/min and consisted of a gradient run using water with 0.1% acetic acid (Solvent A) and acetonitrile isopropanol, 90:10 (Solvent B) for 15 minutes (0-12 min from 25% B to 95% B, 12-12.5 min 95% B, 12.5-15 min 25% B). Electrospray ionization operated in negative ion mode with capillary set at 2.7 kV, desolvation temperature set at 600°C, and source temperature set to 150°C. Optimization of oxylipin MRM transitions were identified by direct injection of pure standards onto the mass spectrometer and using cone voltage and collision energy ramps to optimize detection and most prevalent daughter fragments. Calibration curves were generated prior to each run using standards for each oxylipin. Peak detection and integrations were achieved through Target Lynx (Waters, Milford, MA) and each peak inspected for accuracy and corrected when needed.

#### Statistics

All data are presented as mean ± SEM, and a Shapiro-Wilk test was used to test for a normal distribution. Comparisons of multiple groups were performed using a two-way ANOVA with repeated measures and a Tukey’s post-test with Prism 10.0 (GraphPad Prism, San Diego, CA). P<0.05 was considered significant. Results of post-test comparisons are only shown when the primary interaction was significant.

## Results

### Ffar4 and ChemR23 are co-expressed in adult CMs

Here, we have advanced a hypothetical model for a CM-autonomous signaling mechanism in which Ffar4 activation leads to the production of the ligand for another GPCR, ChemR23 to protect CMs from oxidative stress (Figure 1). As a requisite for this hypothesis to be correct, both Ffar4 and ChemR23 must be co-expressed in CMs. To evaluate this assertion, we created cDNA libraries from individual adult CMs and used RT-qPCR to measure receptor expression. In support of our hypothesis, we detected Ffar4 and ChemR23 co-expression in all the CMs tested (Figure 2).

**Figure 2.**
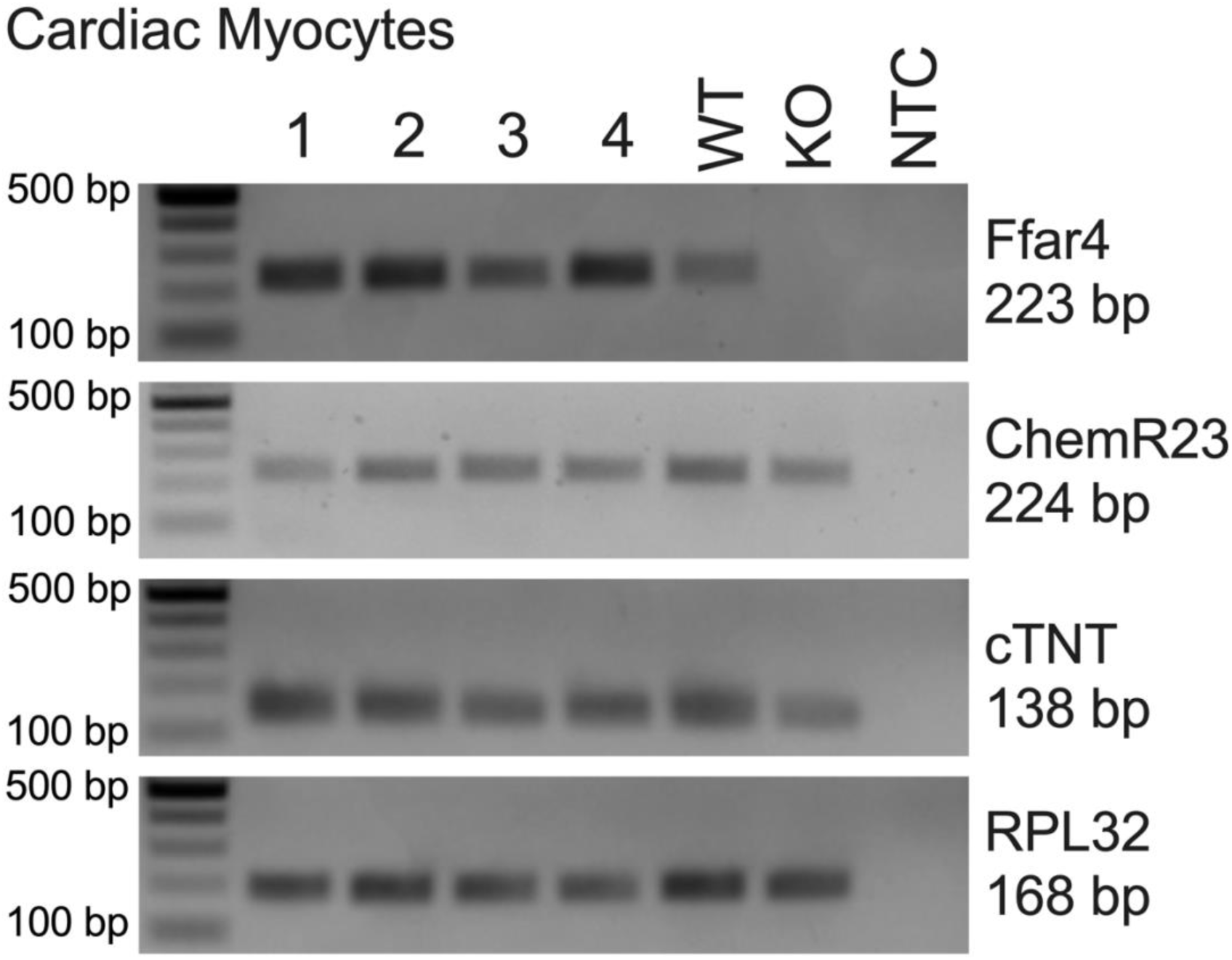
Ffar4 and ChemR23 are co-expressed in adult CMs. Adult cardiac myocytes were isolated from the hearts of WT mice. A cDNA library was generated for each individual CM followed by a pre-amplification procedure as described. Ffar4, ChemR23, cardiac troponin T (cTnT, a CM control), and RPL32 (a library control) were amplified in separate reactions by qRT-PCR using primers and reaction conditions as described. To validate the PCR procedure, reaction products were separated on an agarose gel, and all reactions demonstrate specific amplicons at the predict molecular weight. Lanes 1-4: 4 individual CMs; Lane 5 (WT): whole heart tissue from a WT mouse; Lane 6 (KO): whole heart tissue from an Ffar4KO mouse; Lane 7 (NTC): a non-template control.

### Ffar4 (TUG-891) attenuates cell death and ROS generation induced by hypoxia-reoxygenation (H/R) in adult CMs

We recently demonstrated that CM-specific overexpression of Ffar4 attenuated systolic dysfunction post-I/R (7). To define the mechanistic basis for the purported Ffar4 cardioprotective effects post-I/R, we developed an *in vitro* model of H/R in which adult CMs were subjected to 3 hours of hypoxia followed by 17 hours of reoxygenation to simulate I/R *in vitro*. In adult CMs subjected to this 3/17-H/R protocol, H/R increased CM death by 48.67% ± 7.82% (percent increase in cell death ± SEM) (Figure 3A and 3B, green symbols). Interestingly, the Ffar4-specific agonist, TUG-891 (10 μM), attenuated H/R-induced CM death by 126.09% ± 16.48% (Figure 3A and 3B, closed symbols). Additionally, we measured reactive oxygen species (ROS) generation as a marker of oxidative stress after 3 hours of hypoxia and 4 hours of reoxygenation. The 4 hour reoxygenation timepoint was chosen as a time at which CMs were actively dying. The 3/4-H/R protocol increased CM ROS levels by 85.99% ± 23.27% (Figure 3C and 3D, green symbols), whereas TUG-891 attenuated H/R-induced ROS generation by 117.50% ± 23.00% (Fig. 3C and 3D, closed symbol).

**Figure 3.**
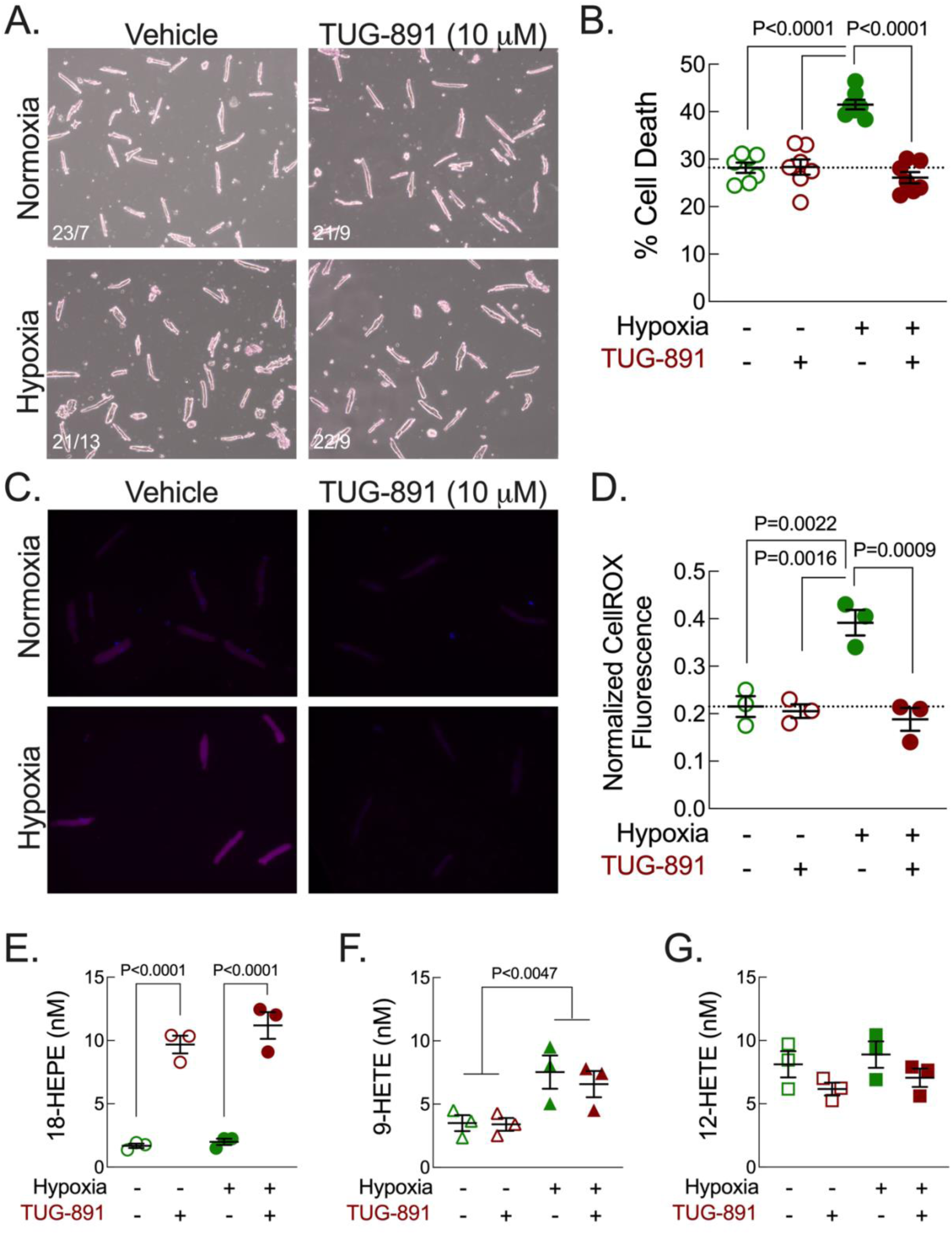
Ffar4 (TUG-891) attenuates cell death and ROS generation induced by hypoxia-reoxygenation in adult CMs. Adult cardiac myocytes were isolated and cultured from the hearts of WT mice, treated overnight with TUG-891 (10 μM) and then subjected to 3 hr of hypoxia followed by 17 hr of reoxygenation. (**A-B**). Cell death was assessed by morphology (rod shape: live; round: dead; (A) CM morphology; (B) quantitation) and (**C-D**) ROS was measured by CellROX fluorescence ((C) CM fluorescence; (D) quantitation). (**E-F**) Quantification of (E) 18-HEPE levels (F) 9-HETE, n=3 (marker of oxidative stress). Data in **B** (n=7), **D** (n=3), **E** (n=3), **F** (n=3), and **G** (n=3) are presented as mean ± SEM and data were analyzed by two-way ANOVA with a Tukey’s multiple comparison post-test to detect differences between groups. For determination of a primary interaction, P˂0.05 was considered significant, and significance between groups (P˂0.05) identified by the Tukey’s post-test are indicated in the figures.

Previously, we observed that activation of Ffar4 in adult CMs induced the production of 18-HEPE (5), whereas loss of Ffar4 reduced 18-HEPE levels in the heart and HDL, which correlated with worsened outcomes in HF (5, 6). Here, we measured Ffar4-mediated production of 18-HEPE in our 3/17-H/R model. In adult CMs, H/R had no effect on 18-HEPE levels, but we confirmed our previous result that TUG-891 (10 μM) increased 18-HEPE levels in adult CMs, regardless of H/R (Figure 3E).

Interestingly, the 3/17-H/R model increased 9-HETE levels, but 9-HETE levels were not affected by TUG-891 (Figure 3F). 9-HETE is produced solely through oxidative stress, indicating the effects of TUG-891 on 18-HEPE were specific, not simply a result of oxidative stress. Finally, TUG-891 showed a trend to decrease the levels of the pro-inflammatory oxylipin 12-HETE regardless of H/R (Figure 3G). This correlates with our previous results indicating that loss of Ffar4 was associated with increased 12-HETE levels in heart and HDL *in vivo* (5, 6), and suggests that Ffar4 attenuates 12-HETE production.

### Figure 4. 18-HEPE and RvE1 attenuate cell death induced by hypoxia-reoxygenation in adult CMs

To test our hypothesis that Ffar4 cardioprotection is due to an Ffar4-18-HEPE-RvE1-ChemR23 signaling pathway, adult CMs were subjected to the 3/17-H/R protocol and treated with 18-HEPE (100 nM, Figure 4A and 4B) or RvE1 (100 nM, Figure 4C and 4D). Importantly, both 18-HEPE and RvE1 attenuated H/R-induced CM death by 99.00% ± 11.07% (Figure 4A and 4B, closed symbols) and 105.31% ± 7.46% (Figure 4C and 4D, closed symbols) respectively. For both 18-HEPE and RvE1, the magnitude of protection was comparable to TUG-891.

**Figure 4.**
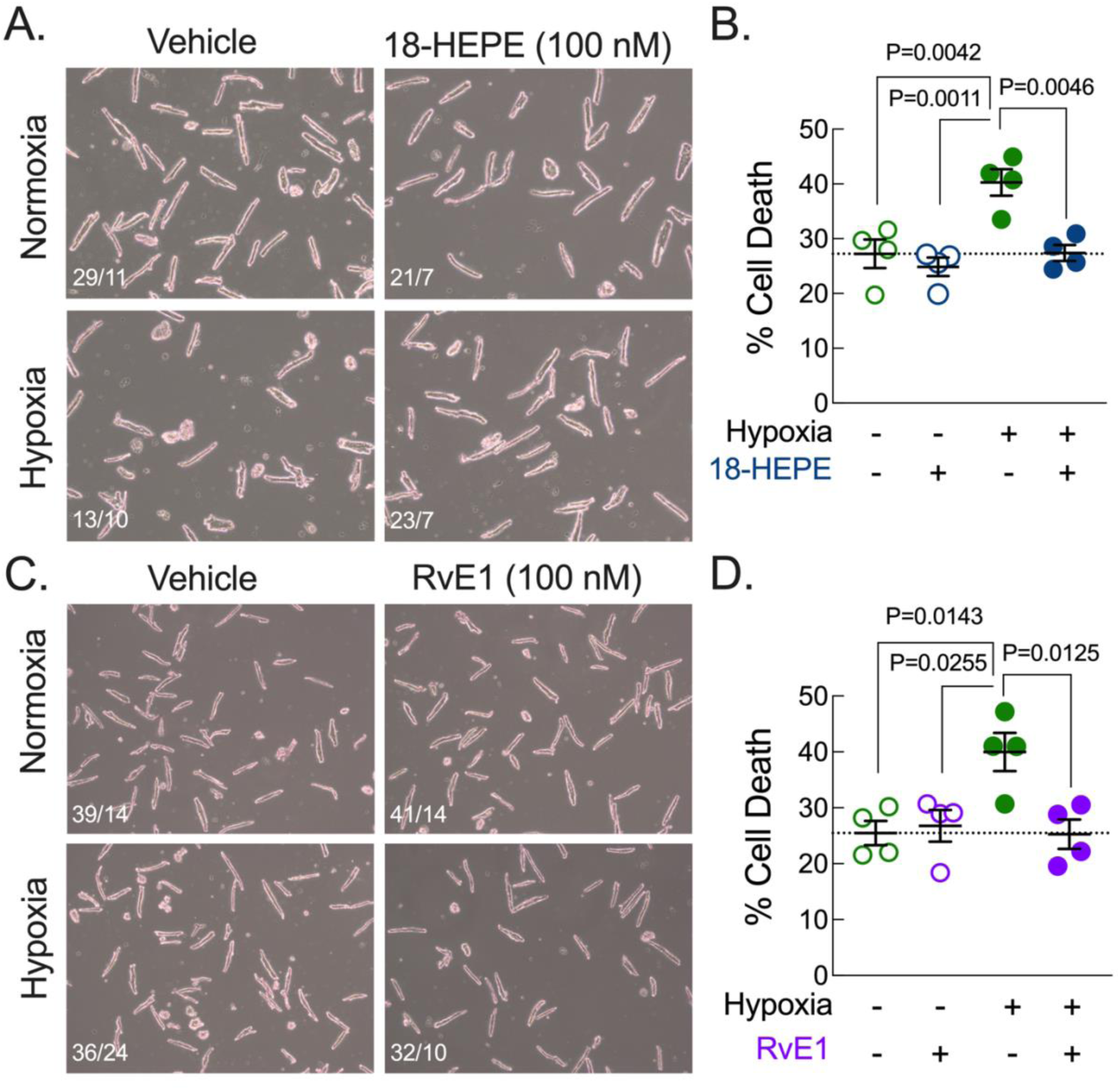
18-HEPE and RvE1 attenuate cell death induced by hypoxia-reoxygenation in adult CMs. Adult cardiac myocytes were isolated and cultured from the hearts of WT mice, treated with **(A-B**) 18-HEPE (100 nM) or (**C-D**) RvE1 (100 nM) for 2 hours and then subjected to 3 hr of hypoxia followed by 17 hr of reoxygenation. (**A, C**) Cell death was assessed by morphology (rod shape: live; round: dead) and (**B, D**) quantified. Data in **B** (n=4) and **D** (n=4) are presented as mean ± SEM and data were analyzed by two-way ANOVA with a Tukey’s multiple comparison post-test to detect differences between groups. For determination of a primary interaction, P˂0.05 was considered significant, and significance between groups (P˂0.05) identified by the Tukey’s post-test are indicated in the figures.

### Figure 5. ChemR23 is required for Ffar4 (TUG-891) attenuation of cell death induced by hypoxia-reoxygenation in adult CMs

To determine if ChemR23 is required for Ffar4 cardioprotection, adult CMs were subjected to the 3/17-H/R protocol and treated with or without TUG-891 (10 μM) and with or without the ChemR23 antagonist α-NETA (100 nM). TUG-891 attenuated H/R-induced CM death (Figure 5A and 5B, green vs maroon symbols), whereas α-NETA ablated TUG-891 cardioprotection by 133.72% ± 14.44% (Figure 5A and 5B, maroon vs blue/maroon symbols). In short, this result indicates that ChemR23 is required for Ffar4 cardioprotection in adult CMs.

**Figure 5.**
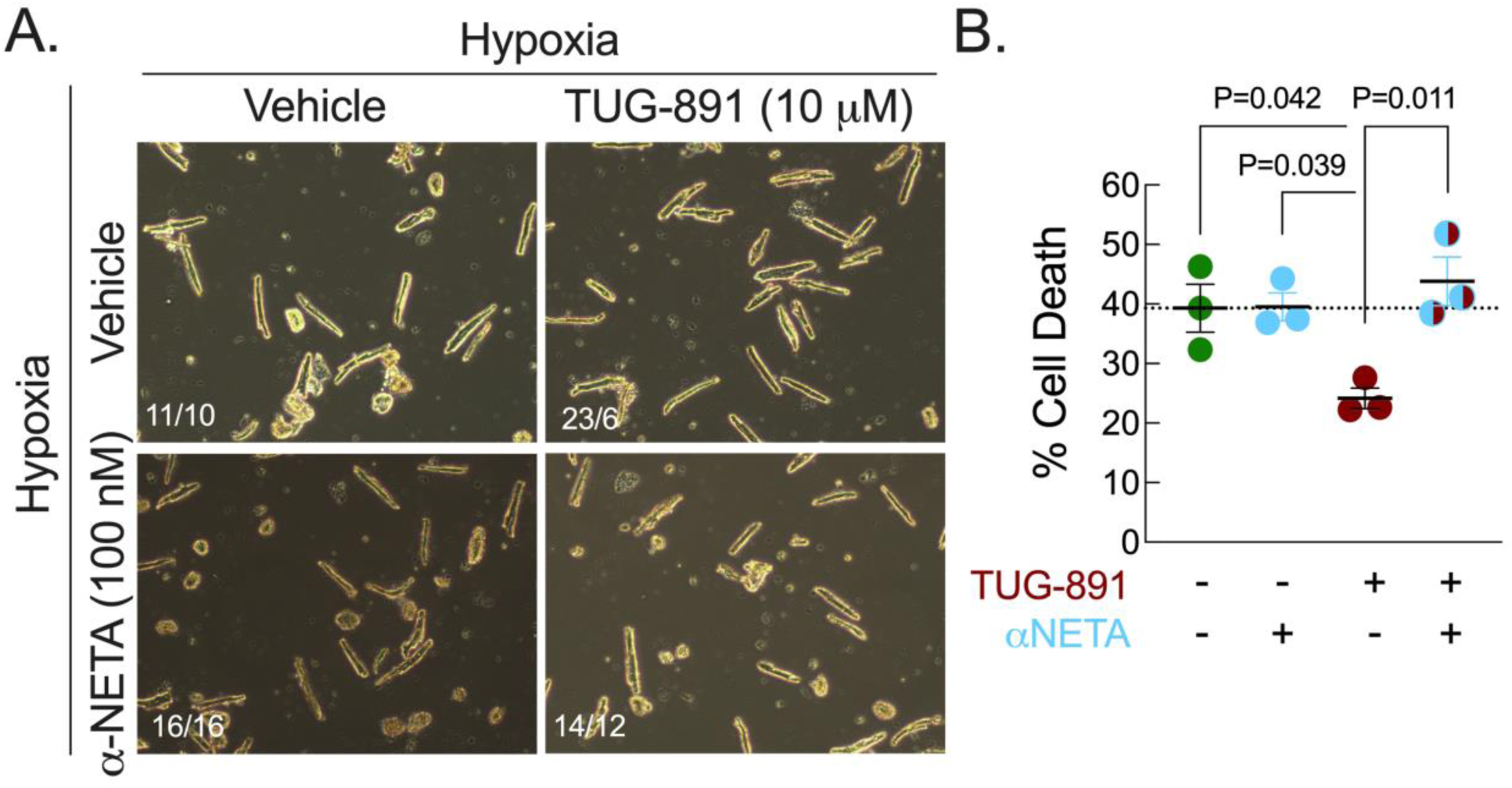
ChemR23 is required for Ffar4 (TUG-891) attenuation of cell death induced by hypoxia-reoxygenation in adult CMs. Adult cardiac myocytes were isolated and cultured from the hearts of WT mice, treated with TUG-891 (10 μM), αNETA (100 nM) or the combination of TUG-891 and αNETA overnight and then subjected to 3 hr of hypoxia followed by 17 hr of reoxygenation. (**A**) Cell death was assessed by morphology (rod shape: live; round: dead) and (**B**) quantified. Data in **B** (n=3) are presented as mean ± SEM and data were analyzed by two-way ANOVA with a Tukey’s multiple comparison post-test to detect differences between groups. For determination of a primary interaction, P˂0.05 was considered significant, and significance between groups (P˂0.05) identified by the Tukey’s post-test are indicated in the figures.

## Discussion

We have proposed the novel hypothesis that a CM intrinsic Ffar4-18-HEPE/RvE1-ChemR23 signaling axis protects CMs from oxidative stress to attenuate ICM as depicted in Fig. 1. Briefly, we show that Ffar4 and ChemR23 are co-expressed in CMs (Fig 2), supporting our proposed cell autonomous signaling pathway. Further, we found that Ffar4 attenuated cell death (Figure 3A and 3B) and oxidative stress (Figure 3C and 3D) in an *in vitro* model of H/R, while similar results were observed with 18-HEPE (Fig 4A and 4B), and RvE1 (Fig 4C and 4D). Lastly, we found that inhibition of ChemR23 prevented TUG-891 induced cardioprotection in H/R (Fig 5). In total, these data support our hypothesis that Ffar4 signaling is cardioprotective via an 18-HEPE/Rve1-ChemR23 signaling axis.

Clinically, increased oxidative stress in CMs following coronary revascularization post-MI drives ICM (20). In CMs, reperfusion and the associated rapid reintroduction of oxygen and nutrients causes oxidative stress, ROS generation, mitochondrial damage, and CM death (3, 20). Recently, in mice with systemic deletion of Ffar4 (Ffar4KO mice), we found that I/R induced a similar and significant decrease in LV systolic function in both male and female WT and Ffar4KO mice after three days, but from that point, systolic function (ejection fraction) recovered to baseline values in WT mice but continued to decrease and remained suppressed in Ffar4KO mice at four weeks (7).

Transcriptome analysis indicated failure to induce gene programs associated with mitochondrial metabolism/function in the infarcted myocardium from Ffar4KO mice relative to the infarcted myocardium from wild-type mice or non-infarcted myocardium from Ffar4KO mice (7). In agreement with these findings, we showed here that Ffar4 attenuates H/R-induced oxidative stress and cell death in adult CMs *in vitro* (Figure 3). CMs have a high energy demand and a high mitochondrial mass. Therefore, damaged mitochondria and increased ROS production might trigger CM death (21, 22).

Improvements in mitochondrial function might explain the reduction in H/R-induced CM death observed here. Others have also observed that Ffar4 attenuates oxidative stress. In adipocytes, TUG-891 stimulated fat oxidation, mitochondrial respiration, and mitochondrial fission in brown adipose tissue (23). Further, activation of Ffar4 with docosahexaenoic acid (DHA, an ω-3 PUFA) improved mitophagy and protected hepatocytes from oxidative stress (24). Lastly, in the heart post-MI, Ffar4 protects the heart from oxidative stress induced arrythmias (25).

We have clearly demonstrated, here and in prior published work, that Ffar4 activation activates cPLA2α and subsequent 18-HEPE production (Fig 3E) (5, 6). Suzumara *et al*. published that 18-HEPE protects cells from oxidative stress and cell death in a model of diabetic retinopathy (26). Here, we show that 18-HEPE treatment protects CMs from H/R induced cell death (Fig 4A and 4B). 18-HEPE is the precursor to the specialized pro-resolving mediator (SPM), RvE1. SPMs are derived from ω3-PUFAs and resolve inflammation (27), and prior studies in the heart indicate that RvE1 attenuates ischemic injury *ex vivo* (9) and *in vivo* (10). Zhang *et al*. has shown that RvE1 protects the heart from doxorubicin induced myocardial apoptosis and oxidative stress (28). Here, we show that treating CMs with RvE1 protects against H/R-induced cell death (Fig 4C and 4D). RvE1 is an endogenous ligand for the GPCR ChemR23, ChemR23 has been best characterized in the context of inflammation. Few studies have focused on the role of ChemR23 in the heart. One study investigating ChemR23 in the context of diabetes driven cognitive decline showed that ChemR23 activation by either of its endogenous ligands (RvE1 or Chemerin) protected mice from oxidative stress and inhibited NLRP3 inflammasome activation via activation of antioxidant pathways (29).

Here, we demonstrated that Ffar4 cardioprotective signaling required ChemR23. In CMs treated with or without TUG-891 and with or without an inhibitor of ChemR23 (α-NETA), we found TUG-891 induced cardioprotection in H/R was abolished when ChemR23 was inhibited (Fig 5). This data supports our hypothesis that this Ffar4-18-HEPE/RvE1-ChemR23 signaling axis is cardioprotective.

Here, we have described a novel, CM-autonomous Ffar4-ChemR23 signaling pathway that protects CMs from oxidative stress, which we hypothesize will protect the heart from ICM post-MI. Ultimately, targeting Ffar4 with synthetic agonists or endogenous fatty acid ligands, for example icosapent ethyl (Vascepa) might be a novel therapeutic strategy to attenuate ICM post-MI.

## Author Contributions

Conceptualization: SJP, GCS, TDO

Methodology: SJP, CLH, ARA, BAH, GCS, TDO

Formal analysis: SJP, BAH, TDO

Investigation: SJP, CLH, BAG, ARA, GCS, TDO

Writing-Original Draft: SJP, TDO

Writing-Review & Editing: SJP, CLH, BAH, GCS, TDO

Visualization: SJP, TDO

Supervision: GCS, TDO

Funding Acquisition: GCS, TDO

